# Growth-Environment Dependent Modulation of *Staphylococcus aureus* Branched-Chain to Straight-Chain Fatty Acid Ratio and Incorporation of Unsaturated Fatty Acids

**DOI:** 10.1101/047324

**Authors:** Suranjana Sen, Seth R. Johnson, Yang Song, Sirisha Sirobhushanam, Ryan Tefft, Craig Gatto, Brian J. Wilkinson

## Abstract

The fatty acid composition of membrane glycerolipids is a major determinant of *Staphylococcus aureus* membrane biophysical properties that impacts key factors in cell physiology including susceptibility to membrane active antimicrobials, pathogenesis, and response to environmental stress. The fatty acids of *S. aureus* are considered to be a mixture of branched-chain fatty acids (BCFAs), which increase membrane fluidity, and straight-chain fatty acids (SCFAs) that decrease it. The balance of BCFAs and SCFAs in strains USA300 and SH1000 was affected considerably by differences in the conventional laboratory medium in which the strains were grown with media such as Mueller-Hinton broth and Luria broth resulting in high BCFAs and low SCFAs, whereas growth in Tryptic Soy Broth and Brain-Heart Infusion broth led to reduction in BCFAs and an increase in SCFAs. Straight-chain unsaturated fatty acids (SCUFAs) were not detected. However, when the organism was grown *ex vivo* in serum, the fatty acid composition was radically different with SCUFAs, which increase membrane fluidity, making up a substantial proportion of the total (<25%) with SCFAs (>37%) and BCFAs (>36%) making up the rest. Staphyloxanthin, an additional major membrane lipid component unique to *S. aureus*, tended to be greater in content in cells with high BCFAs or SCUFAs. Cells with high staphyloxanthin content had a lower membrane fluidity that was attributed to increased production of staphyloxanthin. *S. aureus* saves energy and carbon by utilizing host fatty acids for part of its total fatty acids when growing in serum. The fatty acid composition of *in vitro* grown *S. aureus* is likely to be a poor reflection of the fatty acid composition and biophysical properties of the membrane when the organism is growing in an infection in view of the role of SCUFAs in staphylococcal membrane composition and virulence.

**Funding:** This work was funded in part by grant 1R15AI099977 to Brian Wilkinson and Craig Gatto and grant 1R15GM61583 to Craig Gatto from the National Institutes of Health

## Introduction

*Staphylococcus aureus* is a worldwide significant pathogen in the hospital and the community. Antibiotic resistance has developed in waves [1] such that we now have methicillin-resistant *S. aureus* (MRSA), vancomycin-resistant *S. aureus* (VRSA) and vancomycin-intermediate *S. aureus* (VISA) [2, 3]. Given the threat of multiply antibiotic-resistant *S. aureus*, various aspects of staphylococcal biology including pathogenicity, antibiotic resistance, and physiology are currently being investigated intensively, in part to support the search for novel anti-staphylococcal agents.

The bacterial cytoplasmic membrane forms an essential barrier to the cell and is composed of a glycerolipid bilayer with associated protein molecules, and is a critical determinant of cell physiology. The biophysical properties of the membrane are to a large extent determined by the fatty acyl residues of membrane phospholipids and glycolipids [4, 5]. The lipid acyl chains influence membrane viscosity/fluidity, and impact the ability of bacteria to adapt to changing environments, the passive permeability of hydrophobic molecules, active transport, and the function of membrane-associated proteins [4-6]. Additionally, membrane fatty acid composition has a major influence on bacterial pathogenesis, critical virulence factor expression [7], and broader aspects of bacterial physiology [8].

*S. aureus* membrane fatty acids are generally considered to be a mixture of branched-chain fatty acids (BCFAs) and straight-chain fatty acids (SCFAs) [9-11], and for a comprehensive review of earlier literature see [12]. Typically, *S. aureus* contains about 65% BCFAs and 35% SCFAs. In *S. aureus* the major BCFAs are odd-numbered iso and anteiso fatty acids with one methyl group at the penultimate and antepenultimate positions of the fatty acid chains, respectively (Fig. 1). BCFAs have lower melting points than equivalent SCFAs and cause model phospholipids to have lower phase transition temperatures [13], and disrupt the close packing of fatty acyl chains [14, 15]. The membrane lipid composition of *S. aureus* is further complicated by the presence of staphyloxanthin, a triterpenoid carotenoid with a C30 chain with the chemical name of α-D-glucopyranosyl-1-*O* -(4,4’-diaponeurosporen-4-oate)-6-*O* (12-methyltetradecanoate) [16] (Fig. 1). Staphyloxanthin, as a polar carotenoid, is expected to have a significant influence on membrane properties with the expectation that it rigidifies the membrane [17], and Bramkamp and Lopez [18] have suggested that staphyloxanthin is a critical component of lipid rafts in *S. aureus* incorporating the organizing protein flotillin. Staphyloxanthin has drawn considerable attention in recent years as a possible virulence factor by detoxifying reactive oxygen species produced by phagocytic cells [19, 20], and as a potential target for antistaphylococcal chemotherapy [21].

**Fig 1.**
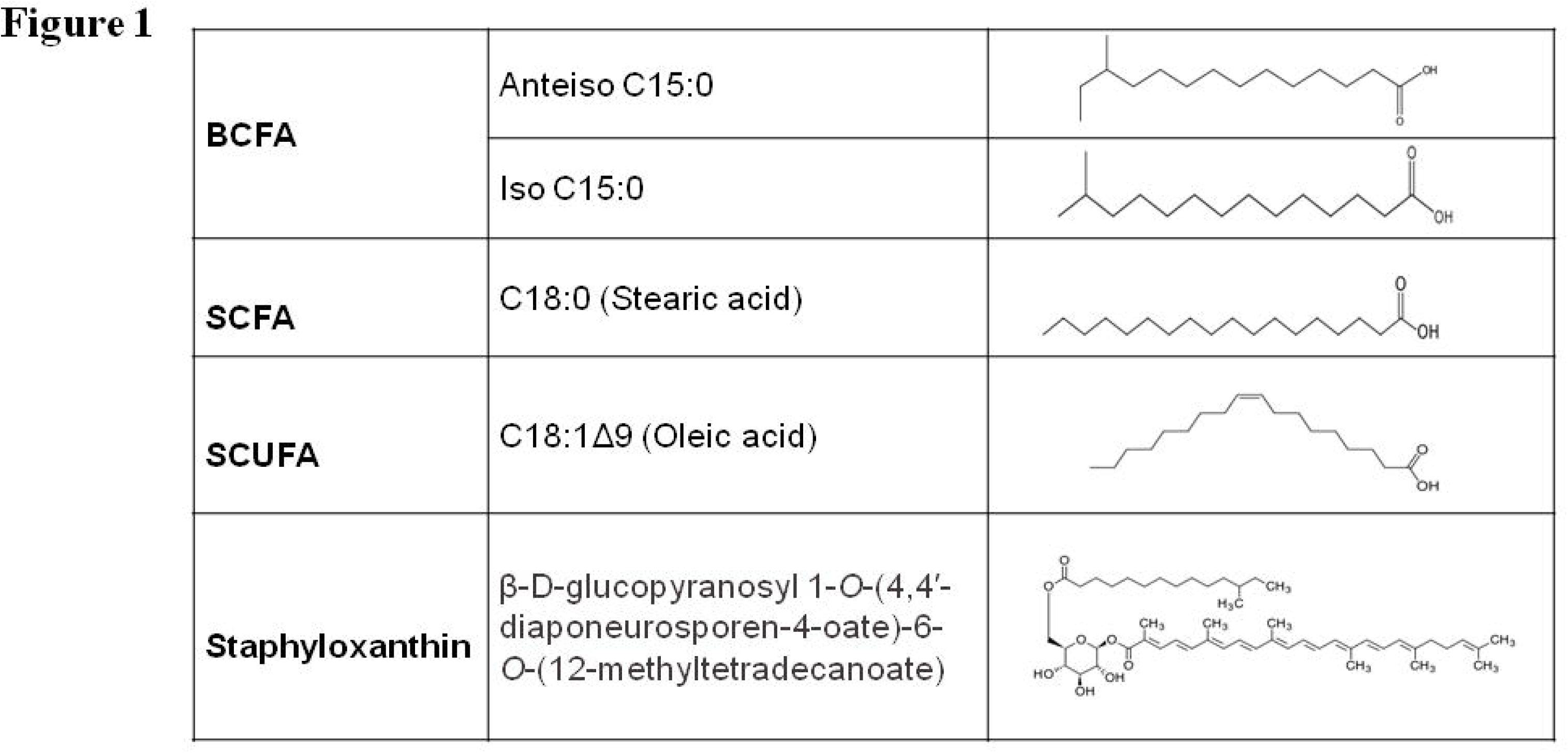
Structures of major fatty acids and staphyloxanthin of the *S. aureus* cell membrane.

In our laboratory, we are interested in the mechanisms of action of and resistance to novel and existing anti-staphylococcal antimicrobials [22-24]. Because much antibiotic work employs Mueller-Hinton (MH) medium, [25] we had occasion to determine the fatty acid composition of a *S. aureus* strain grown in this medium. The analysis was carried out using the MIDI microbial identification system (Sherlock 4.5 microbial identification system; Microbial ID, Newark, DE, USA), [26]. We were taken aback when the fatty acid profile came back showing a very high percentage (84.1%) of BCFAs, and the organism was not even identified by MIDI as a *S. aureus* strain. In a previous study where we grew *S. aureus* in BHI broth we found that 63.5% of the fatty acids were BCFAs, and 32.4% were SCFAs [10]. This is a much more typically observed balance between BCFAs and SCFAs in previous studies of the fatty acid composition of *S. aureus* [9- 12].

A range of different media are used for cultivating *S. aureus* in studies from different laboratories [27]. These are mostly complex media such as TSB, Brain Heart Infusion (BHI) broth, MH broth, Luria-Bertani (LB) broth, and, much more rarely, defined media [11]. Ray et al. [27] and Oogai et al [28] have pointed out that different media have major, but largely unstudied and ignored, effects on the expression of selected target virulence and regulatory genes. Although seemingly prosaic at first glance, issues of choice of strain and medium are nevertheless critical considerations in staphylococcal research [29]. These authors, in their recent protocol publication on the growth and laboratory maintenance of *S. aureus*, have suggested that TSB and BHI media are the media of choice for staphylococcal research. In light of recent literature in various microorganisms, it is becoming glaringly evident that environment has a tremendous effect on the physiology of different pathogens; hence cells from *in vivo* are drastically different from *in vitro* cultured ones. Such distinctions are likely important for studying antimicrobial susceptibilities, drug resistances and pathogenesis.

We decided to carry out a systematic study of the impact of growth medium on the fatty acid and carotenoid composition of *S. aureus* given the large potential impact of these parameters on membrane biophysical properties and its further ramifications. The BCFA: SCFA ratio was significantly impacted by the laboratory medium used, with media such as MH broth encouraging high proportions of BCFAs. However, strikingly, when cells were grown in serum, an *ex vivo* environment, the fatty acid composition changed radically, with straight-chain unsaturated fatty acids (SCUFAs), which were not detected in cells grown in laboratory media, making up a major proportion of the total fatty acids. Biosynthesized bacterial fatty acids are produced by fatty acid synthase II (FASII) [30]. However, fatty acid biosynthesis is expensive in terms of energy and carbon, and many bacteria are able to incorporate extracellular fatty acids into their membrane lipids to varying degrees [30]. This extreme plasticity of *S. aureus* membrane lipid composition is undoubtedly important in determining membrane physical structure and thereby the functional properties of the membrane. The alterations in the fatty acid composition as a result of interactions of the pathogen with the host environment may be a crucial factor in determining its fate in the host. Typically used laboratory media do not result in a *S. aureus* membrane fatty acid composition that closely resembles the likely one of the organism growing *in vivo* in a host.

## Materials and Methods

### Bacterial strains and growth conditions

The primary *S. aureus* strains studied were USA300 and SH1000. USA300 is a community-acquired MRSA strain, and is a leading cause of aggressive cutaneous and systemic infections in the USA [1, 31, 32]. This clinical MRSA also has a well-constructed diverse transposon mutant library [33]. *S. aureu*s strain SH1000, is an 8325-line strain that has been used extensively in genetic and pathogenesis studies [34]. The laboratory media used were MH broth, TSB and Luria Broth (LB) from Difco. For growth and fatty acid composition studies cultures of *S. aureus* strains were grown at 37° C in 250 ml Erlenmeyer flasks containing each of the different laboratory media with a flask–to-medium volume ratio of 6:1. Growth was monitored by measuring the OD_600_ at intervals using a Beckman DU-65 spectrophotometer.

### Growth of *S. aureus* in serum

Sterile fetal bovine serum of research grade was purchased from Atlanta Biologics, USA. The aliquoted serum was incubated in a water bath at 56° C for 30 min to heat inactivate the complement system. *S. aureus* cells were grown for 24 hours in 50 ml of serum in a 250 ml flask at 37°C with shaking at 200 rpm.

### Analysis of the membrane fatty acid composition of *S. aureus* grown in different media

The cells grown in the different laboratory media conventionally used were harvested in mid-exponential phase (OD_600_ 0.6), and after 24 hrs of growth in serum, by centrifugation at 3000 × g at 4° C for 15 minutes and the pellets were washed three times in cold distilled water. The samples were then sent for fatty acid methyl ester (FAME) analysis whereby the fatty acids in the bacterial cells (30-40 mg wet weight) were saponified, methylated, and extracted. The resulting methyl ester mixtures were then separated using an Agilent 5890 dual-tower gas chromatograph and the fatty acyl chains were analyzed and identified by the Midi microbial identification system (Sherlock 4.5 microbial identification system) at Microbial ID, Inc. (Newark, DE) [26]. The percentages of the different fatty acids reported in the tables are the means of the values from three separate batches of cells under each condition. The standard deviations are ±1.5 or less. Some minor fatty acids such as odd-numbered SCFAs are not reported.

### Extraction and estimation of carotenoids

For quantification of the carotenoid pigment in the *S. aureus* cells grown in different media, the warm methanol extraction protocol was followed as described by Davis et al. [35]. Cultures of *S. aureus* were harvested at mid-exponential phase and were washed with cold water. The pellets were then extracted with warm (55°C) methanol for 5 min. The OD_465_ of the supernatant after centrifugation was measured using a Beckman DU 70 spectrophotometer. Determinations were carried out in triplicate.

### Measurement of the fluidity of the *S. aureus* membrane

The fluidity of the cell membrane of the *S. aureus* strains grown in different media were determined by anisotropic measurements using the fluorophore diphenylhexatriene (DPH) following the protocol described previously [36]. Mid exponential phase cells grown in respective media and serum were harvested and washed with cold sterile PBS (pH 7.5).The pellets were then resuspended in PBS containing 2 μM DPH (Sigma, MO) to an OD_600_ of about 0.3 and incubated at room temperature in the dark for 30 min. Fluorescence polarization emitted by the fluorophore was measured using a PTI Model QM-4 Scanning Spectrofluorometer at an excitation wavelength of 360 nm and emission wavelength of 426 nm. The experiments were performed with three separate fresh batches of cells and the Student T-test of the mean polarization values was used to determine statistically significant differences.

## Results

### MH broth and LB increase the content of BCFAs and TSB and BHI broth increase the content of SCFAs

The fatty acid compositions of strain USA300 grown in different laboratory media are shown in Table 1. Growth in MH broth and LB broth resulted in a high content of BCFAs-80.9% and 77.2% respectively, whereas SCFAs were 19.1% and 22.8% respectively. However, in TSB and BHI broth the BCFAs contents were lower at 51.7% and 51.5% respectively, and SCFAs were increased to 48.3 and 48.5% respectively. In MH broth anteiso odd-numbered fatty acids were the major fatty acids in the profile (59.8%), followed by even numbered SCFAs (16.6%), iso odd-numbered fatty acids (15.8%), with iso even-numbered fatty acids making up only a minor portion (4.7%). Anteiso C15:0 was the predominant fatty acid in the membrane lipids (39%). This particular fatty acid has a significant impact on fluidizing membranes [37, 38]. The anteiso fatty acids were significantly reduced in TSB-grown cells (29.3%). The major SCFAs in TSB-grown cells were C18:0 and C20:0 at 19.1% and 18.6% respectively. Overall, the fatty acid compositions were in line with many previous studies of *S. aureus* fatty acid composition [9-12], but we are unaware of previous studies that have identified this impact of medium on the proportions of BCFAs and SCFAs in the membrane.

**Table 1.** The membrane fatty acid profile of *S. aureus* USA300. % (wt/wt) of total fatty acids

| *Growth Medium* | *Anteiso odd* |  |  |  | *Iso odd* |  |  |  | *Iso even* |  |  |  | *Straight Even* |  |  |  |  | *BC FA* | *SC FA* |
| --- | --- | --- | --- | --- | --- | --- | --- | --- | --- | --- | --- | --- | --- | --- | --- | --- | --- | --- | --- |
|  | C15:0 | C17:0 | C19:0 | SUM | C15:0 | C17:0 | C19:0 | SUM | C14:0 | C16:0 | C18:0 | SUM | C14:0 | C16:0 | C18:0 | C20:0 | SUM |  |  |
| **BHI** | 28.5 | 2.6 | ND | 31.1 | 12.2 | 1.8 | ND | 14 | 2.9 | 1.4 | ND | 4.3 | 3.3 | 7.9 | 21.5 | 13.4 | 46.1 | 51.5 | 48.5 |
| **TSB** | 26.9 | 2.4 | ND | 29.3 | 12.8 | 2 | ND | 14.8 | 3.7 | 1.8 | ND | 5.5 | 2.3 | 6.4 | 19.1 | 18.6 | 46.4 | 51.7 | 48.3 |
| **MHB** | 39 | 15 | 5.8 | 59.8 | 7.7 | 4.9 | 3.2 | 15.8 | 1 | 2.1 | 1.6 | 4.7 | 1.1 | 1.8 | 6.5 | 7.2 | 16.6 | 80.9 | 19.1 |
| **LB** | 36.5 | 11 | 2.4 | 49.9 | 13.3 | 6.6 | 2.6 | 22.5 | 1.4 | 2.2 | 1 | 4.6 | ND | 3 | 10 | 6.9 | 19.9 | 77.2 | 22.8 |

ND- Not detected

The results of a similar series of experiments with strain SH1000 are shown in Table 2. Overall, the proportion of BCFAs of this strain was higher than strain USA300. In strain SH1000 the BCFAs were higher than USA300 in all media-BHI 66.6%, TSB 68.5%, with particularly high contents in MH broth 90.2% and LB 89%. The proportion of SCFAs was correspondingly smaller in all cases compared to strain USA300. Anteiso fatty acids were the major class of fatty acids in all media, amongst which anteiso C15:0 was present in the highest amount in all cases. However, the same phenomenon was noted where MH broth and LB encouraged a high proportion of BCFAs, low SCFAs, and TSB and BHI had the opposite effects on fatty acid composition. Two additional media were studied with this strain. Both Tryptone broth [39] and defined medium [40] resulted in high BCFAs (80.4% and 85% respectively), and low SCFAs (19.7% and 15% respectively).

**Table 2.**
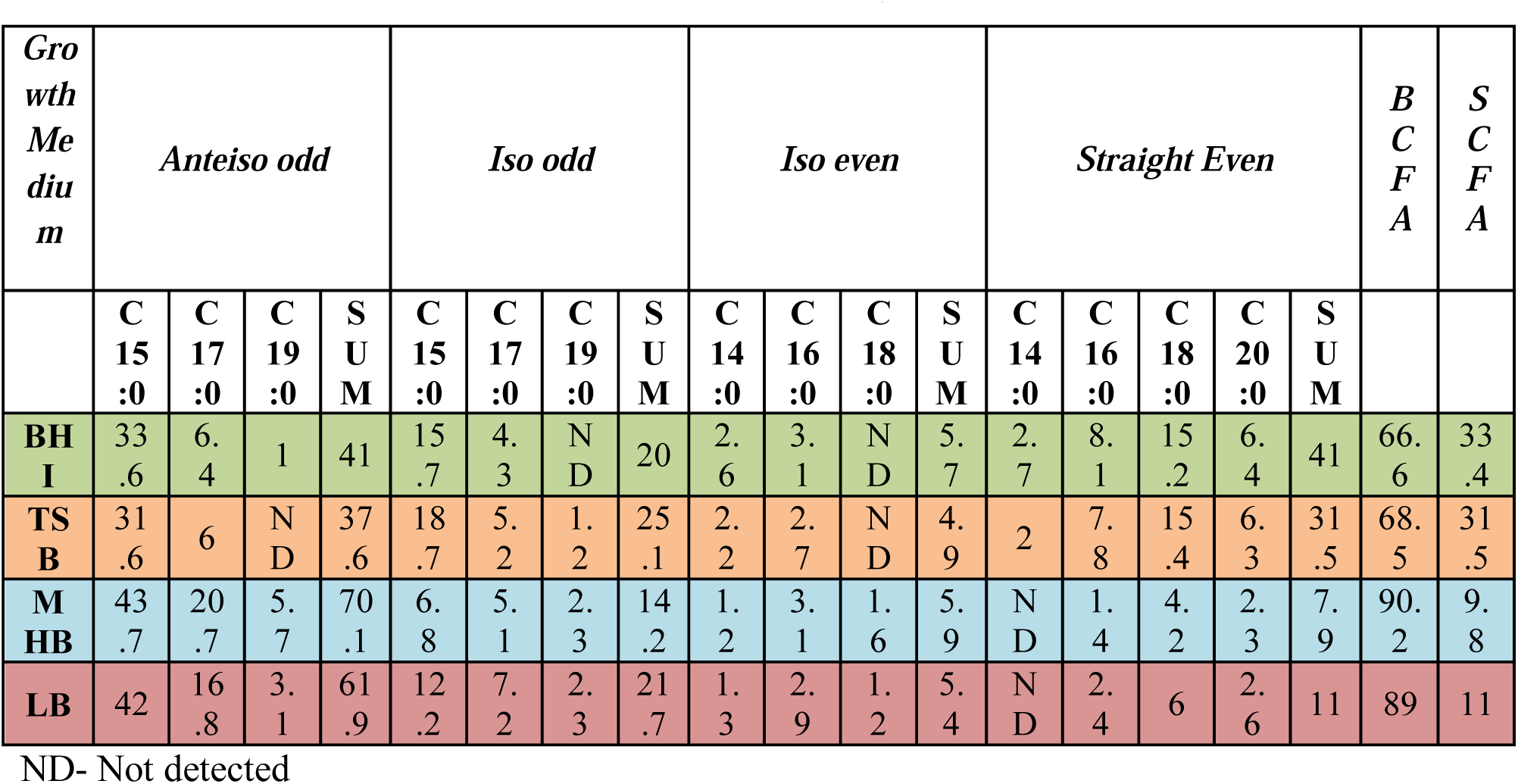
The membrane fatty acid composition of *S. aureus* strain SH1000. % (wt/wt) of total fatty acid

The phenomenon of higher BCFAs in MH broth-grown cells and higher SCFAs in TSB grown cells was observed in 8 out of 9 strains tested which included MRSA, VISA and daptomycin decreased susceptibility strains (data not shown). This indicates that the phenomenon noted in strains USA300 and SH1000 also extends to other *S. aureus* strains.

### The fatty acid composition of *S. aureus* grown *ex vivo* in serum is radically different to those of the organism grown in laboratory media

It was of interest to try and get an idea of the fatty acid composition of *S. aureus* grown in vivo. Strain USA300 and SH1000 were grown in serum, which resulted in major changes in the fatty acid profile (Table 3). Total BCFAs were reduced to 37.5% in USA300 and 36.3 in SH1000; SCFAs were at 37.8% in USA300 and 32.1% in SH1000, but 25% of the fatty acid profile in the case of USA300 and 30.6% in SH1000 was accounted for by SCUFAs. Strikingly, this type of fatty acid was not present in the profile of the organism when grown in laboratory media. Interestingly, BCFAs and SCUFAs have similar effects in increasing fluidity of the membrane [4].

**Table 1.**
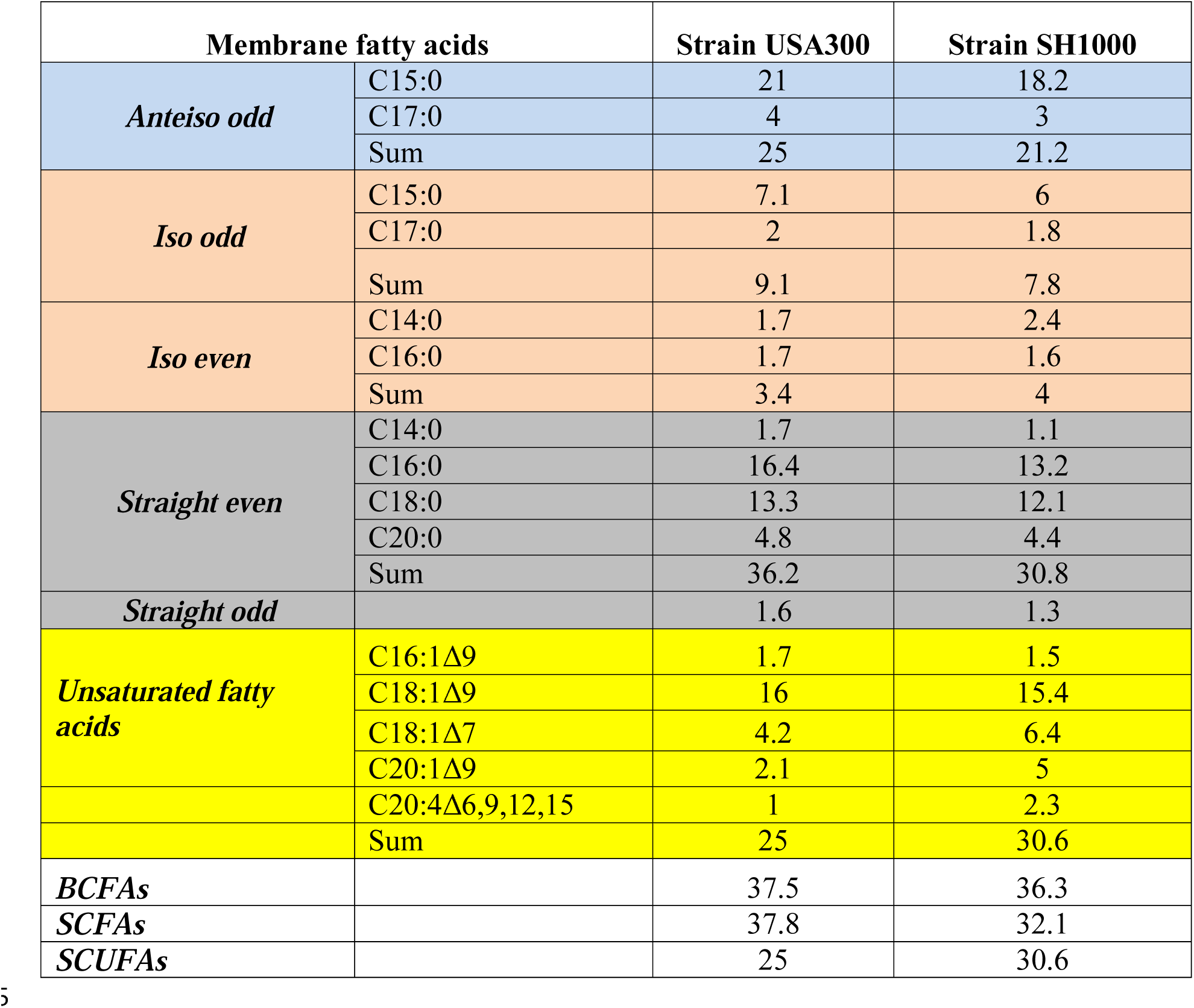
The membrane fatty acid compositions of *S. aureus* USA300 and SH1000 grown *ex vivo* in serum. % (wt/wt) of total fatty acid

### Carotenoid content of cells grown in different media

Staphyloxanthin is another significant membrane component that might impact the biophysical properties of the membrane. Accordingly, the carotenoid content of cells grown in different media were determined and the results are shown in Fig. 2. Strain SH1000 cells grown in MH broth had a much higher carotenoid content than cells grown in the other media. The pellets of cells grown in this particular media were noticeably yellow. It is possible that the carotenoid content rises to counterbalance the potentially high fluidity of MH broth-grown cells with their high content of BCFAs, specifically mainly anteiso fatty acids. LB (high BCFAs) and serum (high SCUFAs) - grown cells had higher carotenoid contents than TSB or BHI broth–grown cells. In strain USA300 MHB-and serum-grown cells also had higher carotenoid contents than did cells grown in BHI, TSB or LB. In general this strain was less pigmented than strain SH1000.

**Fig 2.**
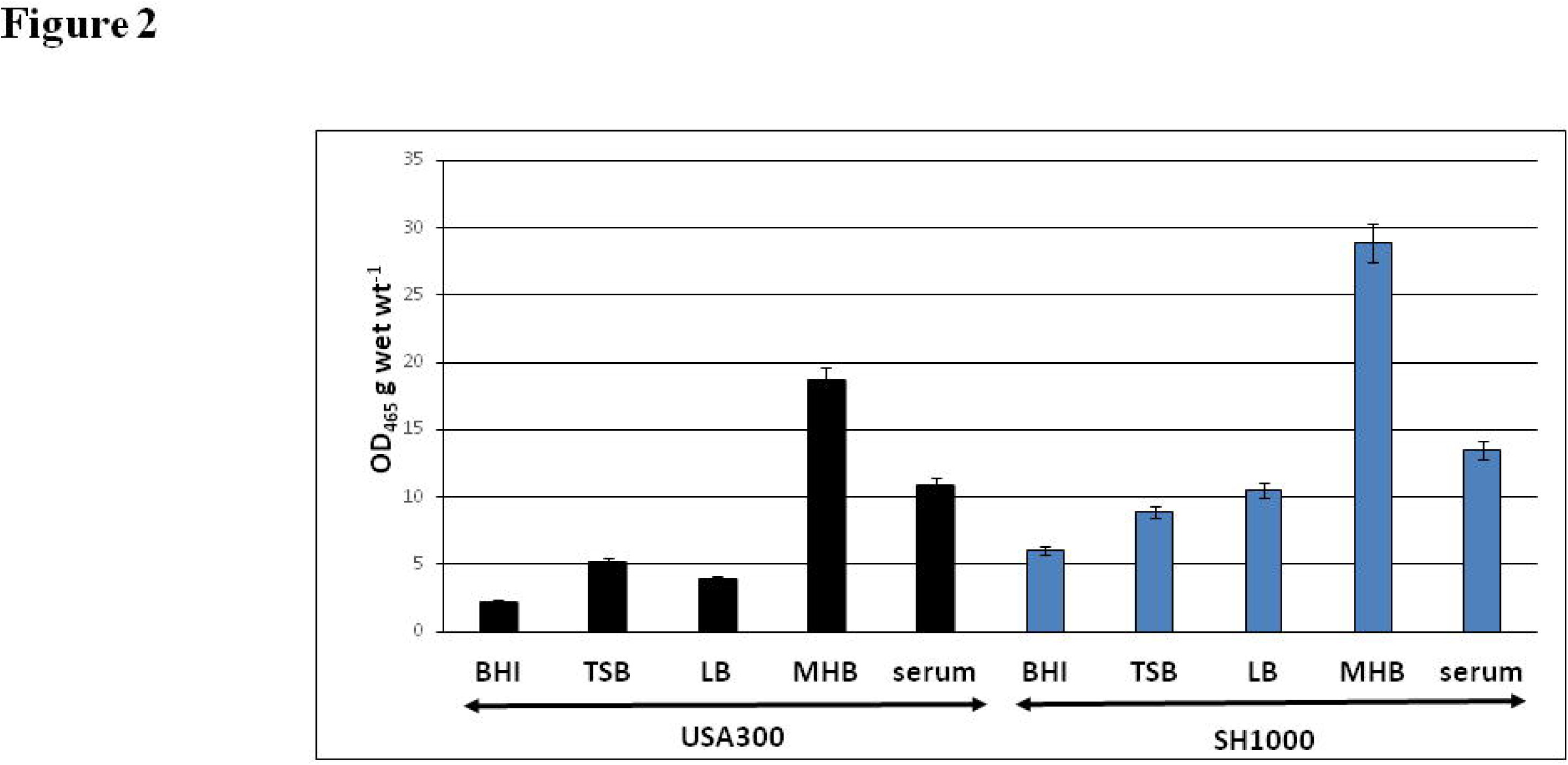
Influence of growth environment on the carotenoid content of *S. aureus*. The strains, USA300 (black columns) and SH1000 (blue columns), were grown in different growth media and the carotenoid was estimated after extraction by warm methanol.

### Membrane fluidity of *S. aureus* cells with different fatty acid compositions

The membrane fluidity of cells of strain SH1000 grown in BHI broth, LB and TSB were very similar (0.185-0.19) as shown in Fig. 3. The membranes of MH-broth and serum-grown cells, 0.25 and 0.248 were significantly less fluid than cells grown in the other media. Possibly the higher carotenoid contents of cells grown in MH broth and serum rigidifies the membrane. Strain USA300 also showed a similar pattern of membrane fluidity in the different growth media Fig. 3. The membrane fluidity of both strains was highest in cells grown in LB, consistent with the high content of BCFAs. Furthermore, there was no accompanying increase in staphyloxanthin content with its possible membrane rigidifying effect in contrast to MHB or serum-grown cells.

**Fig 3.**
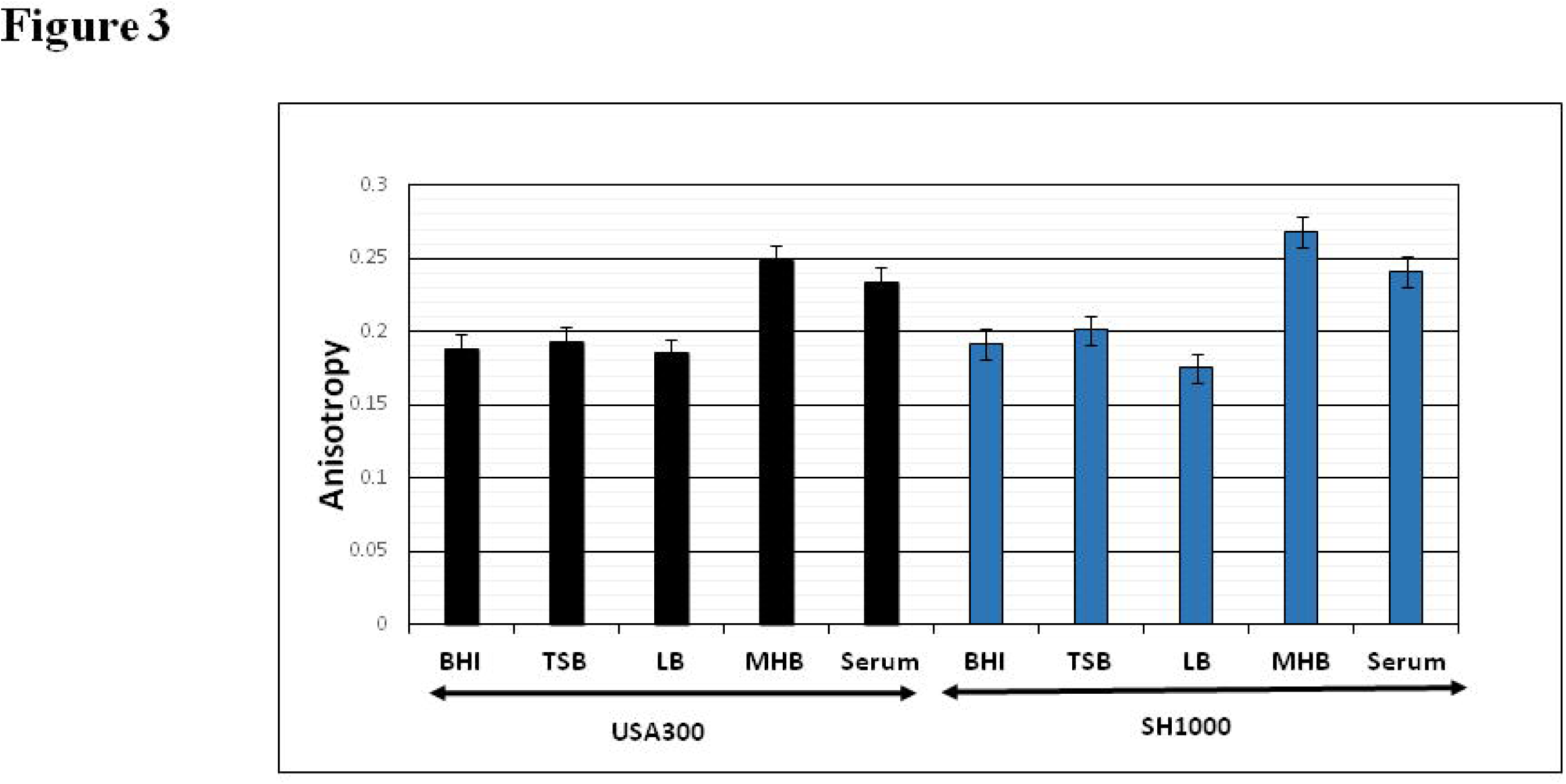
Influence of growth environment on the membrane fluidity of *S. aureus* cells. The strains, USA300 (black columns) and SH1000 (blue columns), were grown in the different media to mid exponential phase and membrane anisotropy was measured by fluorescence polarization.

## Discussion

From numerous studies over the past several decades of *S. aureus* grown *in vitro* in various laboratory media it is considered that the membrane fatty acid composition of the organism is a mixture of BCFA and SCFAs [9-12], and BCFAs have generally been found to be predominant. Through study of a range of different conventional growth media, certain media were found to encourage a higher proportion of BCFAs than others, whereas in some media the proportion of SCFAs was increased. This may have significant physiological ramifications given the opposing effects of BCFAs and SCFAs on membrane fluidity with BCFAs fluidizing and SCFAs rigidifying the membrane [4]. However, there was a radical change in the entire fatty acid composition when the organism was grown *ex vivo* in serum with SCUFAs appearing in the profile in significant amounts accompanied with a decrease in BCFA content.

It is useful to discuss our fatty acid compositional data in the context of what is known about phospholipid biosynthesis and the positional distribution of fatty acids on the 1 and 2 carbon atoms of the glycerol residue (Fig. 4). Phosphatidic acid is a key intermediate in the biosynthesis of the *S. aureus* phospholipids, which are phosphatidyl glycerol, cardiolipin and lysyl-phosphatidyl glycerol [5]. Our current knowledge of the pathway of phospholipid biosynthesis and the incorporation of exogenous and endogenous fatty acids is summarized in Fig. 4 [41]. Phosphatidic acid (PtdOH), the universal precursor of phospholipids, is synthesized by the stepwise acylation of *sn*-glycerol-3-phosphate first by PlsY that transfers a fatty acid to the 1-position from acyl phosphate. The 2-position is then acylated by PlsC utilizing acyl-ACP. Acyl-ACP is produced by the FASII pathway and PlsX catalyses the interconversion of acyl-ACP and acyl phosphate. Exogenous fatty acids readily penetrate the membrane and are activated by a fatty acid kinase to produce acyl phosphate that can be
utilized by PlsY, or they can be converted to acyl-ACP for incorporation into the 2-position by PlsC. Exogenous fatty acids can also be elongated by the FASII pathway. When *S. aureus* is grown in medium that results in a high proportion of BCFAs the major phospholipid, phosphatidyl glycerol (PtdGro), has, almost exclusively, anteiso C17:0 at position 1 and anteiso C15:0 at position 2 [42]. Growth in the presence of oleic acid (C18:1Δ9) showed anteiso C17:0 at position 1 was replaced by C18:1Δ9 and C20:1Δ11, whereas the anteiso C15:0 at position 2 remained at about 50%. BCFAs are not present in serum and hence must be biosynthesized from 2-methylbutyryl CoA, most likely produced from isoleucine.

**Fig 4.**
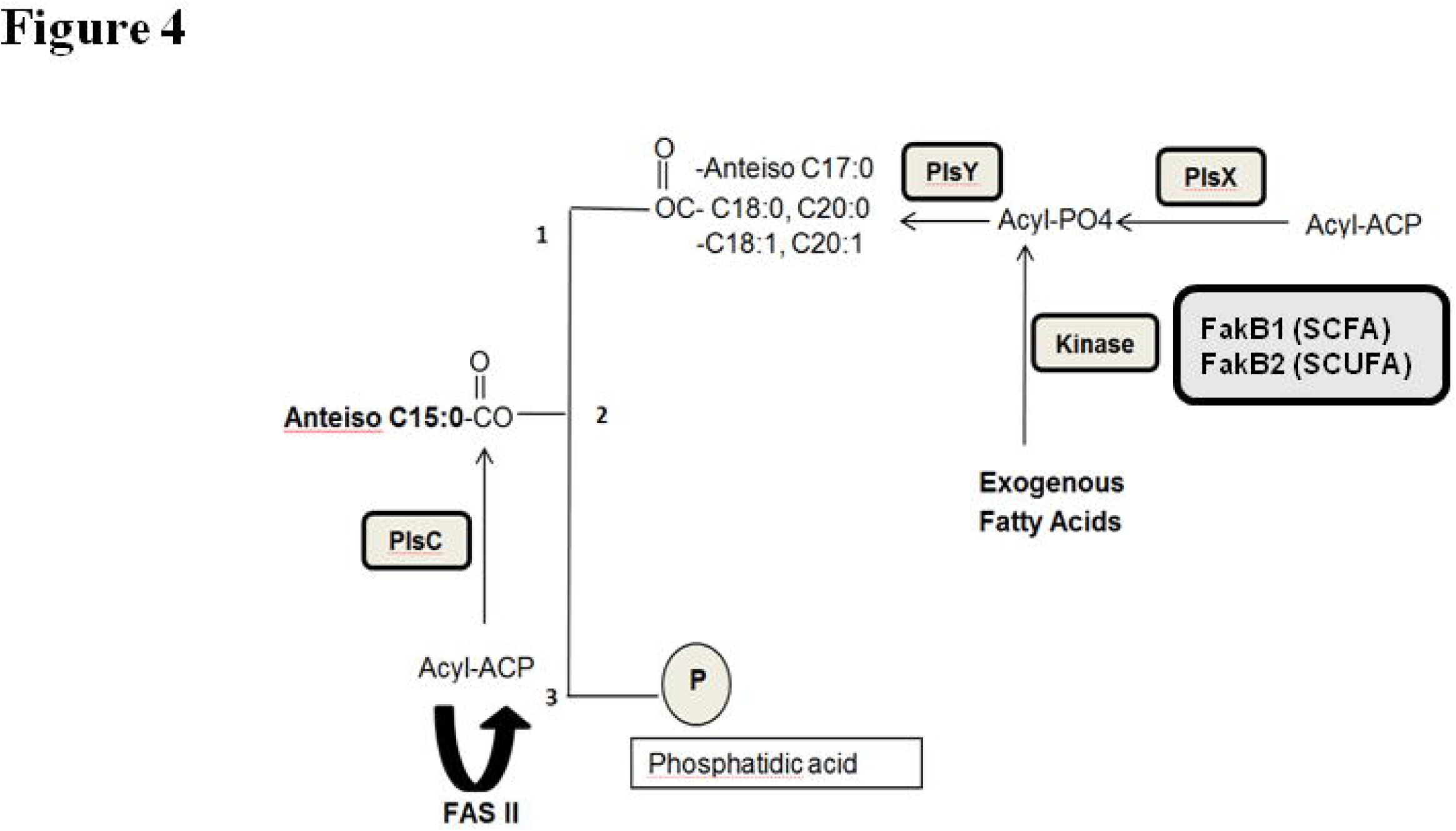
Pathway of phospholipid biosynthesis and the incorporation of exogenous and endogenous fatty acids in *S. aureus*. Phosphatidic acid (PtdOH), the universal precursor of phospholipids, is synthesized by the stepwise acylation of *sn*-glycerol-3-phosphate first by PlsY that transfers a fatty acid to the 1-position from acyl phosphate. The 2-position is then acylated by PlsC utilizing acyl-ACP. Acyl-ACP is produced by the FASII pathway and PlsX catalyses the interconversion of acyl-ACP and acyl phosphate. Exogenous fatty acids readily penetrate the membrane and are activated by a fatty acid kinase ( FakB1 for SCFAs and FakB2 for SCUFAs) to produce acyl phosphate that can be utilized by PlsY, or that can be converted to acyl-ACP for incorporation into the 2-position by PlsC. Exogenous fatty acids can also be elongated by the FASII pathway. Figure modified from Parsons et al. [41].

### What determines the balance between BCFAs and SCFAs in cells grown in laboratory media?

MH medium leads to high proportion of BCFAs in the staphylococcal cells whereas growth in TSB leads to an increase in the proportion of SCFAs. MH broth (Difco) is composed of beef extract powder (2 g/l), acid digest of caseine (17.5 g/l), and soluble starch (1.5 g/l). Thus, by far the major medium component is acid digest of caseine, and this is expected to be high in free amino acids. TSB (Difco) is composed of pancreatic digest of caseine (17 g/l), enzymatic digest of soybean meal (3 g/l), dextrose (2.5 g/l), sodium chloride (5 g/l) and dipotassium phosphate (2.5 g/l). The major components then of TSB are a mixture of peptides formed by enzymatic digestion of caseine and soybean meal. Payne and Gilvarg [43] fractionated Bacto Neopeptone using gel filtration. They found that peptides with a molecular weight below 650 represented about 25% of the mixture, and free amino acids were about 1 % of the entire preparation. We believe that the free amino acids from the acid digest of casein can have a dominant effect on the fatty acid composition.

*Listeria monocytogenes* is a Gram-positive bacterium with a very high (90%) proportion of BCFAs in its cell membrane. The fatty acid composition of the organism grown in defined medium not containing any branched-chain amino acids was readily modified by exogenous isoleucine or leucine, which resulted in the fatty acid profile being dominated by anteiso odd and iso odd fatty acids respectively [26]. *L. monocytogenes* can also obtain amino acids by metabolism of peptides that are taken up [44] as can *S. aureus* [45]. It may be that transport of free branched-chain amino acids results in higher pool levels than when they are biosynthesized or produced through metabolism of transported peptides, giving them a dominant effect on fatty acid composition. In *S. aureus* supplementation of medium with 2-methylbutyrate, a precursor of anteiso fatty acids, significantly increased the content of anteiso C15:0 and C17:0 [10]. Mutants of *S. aureus* in the transporters of leucine and valine lacked odd and even numbered fatty acids derived from these amino acids [46].

Growth in media such as TSB and BHI lead to a higher proportions of SCFAs than media such as MH broth, although SCUFAs were not detected. The origin of SCFAs is not clear as to whether they originate from the medium or are biosynthesized. Typically in bacteria SCFAs are biosynthesized from acetyl CoA via the activities of FabH. However, acetyl CoA was a poor substrate for *S. aureus* FabH [47], whereas the enzyme had high activity for butyryl CoA raising the possibility that butyrate is the primer for biosynthesis of SCFAs in *S. aureus*. It is also possible that SCFAs that may be present in TSB and BHI may be utilized directly for fatty acid elongation to the SCFAs in the membrane typical of growth in these media.

### The underappreciated ability of S. aureus to incorporate host fatty acids from serum

A striking finding in our paper is that *S. aureus* has the capacity to incorporate large proportions of SCFAs and SCUFAs when grown *ex vivo* in serum. Earlier reports of *S. aureus* fatty acid composition have not reported significant amounts of SCUFAs in *S. aureus* [9-12]. Indeed it appears that *S. aureus* lacks the genes necessary to biosynthesize unsaturated fatty acids [41]. However an early report by Altenbern [48] showed that inhibition of growth by the fatty acid biosynthesis inhibitor cerulenin could be relieved by SCFAs or SCUFAs, implying *S. aureus* had the ability to incorporate preformed fatty acids. Fatty acid compositional studies of the cells were not reported though. Serum is lipid rich [49-51] and a comprehensive analysis of the human serum metabolome including lipids has recently been published [52]. BCFAs are present, if at all, in only very small amounts in serum. Bacterial pathogens typically have the ability to incorporate host-derived fatty acids thereby saving carbon and energy since fatty acids account for 95% of the energy requirement of phospholipid biosynthesis [30].

The FASII pathway has been considered to be a promising pathway for inhibition with antimicrobial drugs. The viability of FASII as a target for drug development was challenged by Brinster et al. [53] especially for bacteria such as streptococci where all the lipid fatty acids could be replaced by SCFAs and SCUFAs from serum. However, Parsons et al. [42] showed that exogenous fatty acids could only replace about 50% of the phospholipid fatty acids in *S. aureus* and concluded that FASII remained a viable drug target in this organism.

The relationship between *S. aureus* and long-chain SCUFAs and SCFAs is a complex one. On one hand these fatty acids in the skin and other tissues form part of the innate defense system of the host due to their antimicrobial activities [54-56]. Very closely related structures can either be inhibitory to growth at low concentrations, or can have little effect on growth at relatively high concentrations [39, 57-59]. For example C16:1Δ6 and C16:1Δ9 are highly inhibitory whereas C18:1Δ9 and C18 are not inhibitory and are actually incorporated into the phospholipids by this pathogen [39].

The enzyme fatty acid kinase (Fak) responsible for incorporation of extracellular fatty acids into *S. aureus* phospholipids [41], is also a critical regulator of virulence factor expression [60], and biofilm formation [61]. Fak phosphorylates extracellular fatty acids for incorporation into *S. aureus* membrane phospholipids [41]. FakA is a protein with an ATP-binding domain that interacts with FakB1 and FakB2 proteins that bind SCFAs and SCUFAs preferentially respectively. Fatty acid kinase activity producing FakB (acyl-PO_4_) was proposed to be involved in the control of virulence gene expression. Interestingly FakB2 shows a high degree of specificity for C18:1Δ9, a fatty acid not produced by *S. aureus*, and may act as a sensor for the host environment via the abundant mammalian fatty acid C18:1Δ9 [41], which is subsequently incorporated into the membrane lipids.

Besides occurring in membrane phospholipids and glycolipids fatty acids are present in lipoproteins at their N terminus in the form of an N-acyl-S-diacyl-glycerol cysteine residue and an additional acyl group amide linked to the cysteine amino group [62]. It is estimated that there are 50-70 lipoproteins in S. *aureus*, many of them involved in nutrient acquisition. Additionally, lipoproteins contribute important microbe-associated molecular patterns that bind to Toll-like receptors and activate innate host defense mechanisms. Recently, Nguyen et al. [63] have shown that when *S. aureus* is fed SCUFAs they are incorporated into lipoproteins and the cells have an increased toll-like receptor 2- dependent immune stimulating activity, which enhances recognition by the immune defense system.

### Changes in staphyloxanthin in cells grown under different conditions with different membrane fatty acid compositions

The carotenoid staphyloxanthin is a unique *S. aureus* membrane component that affects membrane permeability, defense against reactive oxygen species, and is a potential drug target. It appeared that cells grown in media encouraging a high proportion of BCFAs or in serum resulting in high SCUFAs, both of which would be expected to increase membrane fluidity, tended to have higher staphyloxanthin contents. However, despite having high amounts of fluidizing BCFAs or SCUFAs, cells grown in MH broth or serum had cellular membranes that were significantly less fluid. Thus it may be inferred that the pigment staphyloxanthin could actually be preventing the membrane from becoming hyper fluid under the particular growth conditions which yield an unusually high amounts of BCFAs or SCUFAs. These conditions may thus result in staphylococcal cells which have a better chance at surviving against oxidative stress and host defense peptides [20]. However this relationship is likely to be complex in that LB-grown cells that had high BCFAs did not have high carotenoid levels, and the phenomenon deserving of more detailed investigation. Interestingly, in the biosynthesis of staphyloxanthin, the end step involves an esterification of the glucose moiety with the carboxyl group of anteiso C15:0 by the activity of the enzyme acyltransferase CrtO [16]. Thus the availability of specific fatty acid precursors and the lipid metabolism may play a significant role in pigment production.

### Plasticity of *S. aureus* membrane lipid composition and its possible ramifications in membrane biophysics and virulence

Given the crucial role of the biophysics of the membrane in all aspects of cell physiology, such radical changes in the membrane lipid profile can have significant but as yet undocumented impacts on critical functional properties of cells such as virulence factor production, susceptibilities to antimicrobials and tolerance of host defenses. It is important to assess the biophysical and functional properties of the membranes of the cells with such radically different fatty acid compositions. Although BCFAs and SCUFAs both increase membrane fluidity, they do not yield cells with identical morphologies [14], or fitness for tolerating cold stress [64]. Also a *S. aureus* fatty acid auxotroph created by inactivation of acetyl coenzyme A carboxylase (*ΔaccD*) was not able to proliferate in mice, where it would have access to SCFAs and SCUFAs [65]. Due to the ability of a pathogen to adapt and undergo dramatic alterations when subjected to a host environment, there is a growing appreciation in the research community for the fact that the properties of the organism grown *in vivo* are probably very different from when it is grown *in vitro*. This distinction may have a huge impact on critical cellular attributes controlling pathogenesis and resistance to antibiotics. Expression of virulence factors is significantly different in serum-grown organisms [28], and there are global changes in gene expression when *S. aureus* is grown in blood [66]. *S. aureus* grown in serum or blood will have different membrane lipid compositions than cells grown in laboratory media and this may have a significant impact on the expression of virulence factors and pathogenesis of the organism.

We have demonstrated a hitherto poorly recognized growth environment-dependent plasticity of *S. aureus* membrane lipid composition. The balance of BCFAs and SCFAs was affected significantly by the variations in laboratory medium in which the organism grew. SCUFAs became a major membrane fatty acid component when the organism was grown in serum. These findings speak to the properties of pathogens grown *in vitro* versus *in vivo*. In 1960 Garber [67] considered the host as the growth medium and the importance of the properties of the pathogen at the site of infection. There has been a renewed appreciation of this in recent years [68]. Massey et al. [69] showed that *S. aureus* grown in peritoneal dialysate acquired a protein coat. Krismer et al. [70] devised a synthetic nasal secretion medium for growth of *S. aureus*. Tn-seq analysis has been used to identify genes essential for survival in infection models versus rich medium. Citterio et al. [71] reported that the activities of antimicrobial peptides and antibiotics were enhanced against various pathogenic bacteria by supplementation of the media with blood plasma to mimic in vivo conditions. In order to replicate a membrane fatty acid composition more closely resembling that of the bacteria growing *in vivo*, it may be desirable to supplement laboratory media with SCFAs and SCUFAs.

## References

1. Chambers HF, and Deleo FR. Waves of resistance: Staphylococcus aureus in the antibiotic era. Nat Rev Microbiol. 2009; 7: 629–641. PMID: 19680247

2. Howden BP, Davies JK, Johnson PD, Sinear TP, and Grayson ML. Reduced vancomycin susceptibility in Staphylococcus aureus, including vancomycin-intermediate and heterogeneous vancomycin-intermediate strains: resistance mechanism, laboratory detection, and clinical implications. Clin Microbiol. Rev. 2010; 23: 99–139. PMID: 20065327

3. Klevens RM, Morrison MA, Nadle J, Petit S, Gershman K, et al. Invasive methicillin-resistant Staphylococcus aureus infections in the United States. JAMA. 2007; 298:1763–1771. PMID: 17940231

4. Zhang YM, and Rock CO. Membrane lipid homeostasis in bacteria. Nat Rev Microbiol. 2008; 6: 222–233. PMID: 18264115.

5. Parsons JB, and Rock CO. Bacterial lipids: metabolism and membrane homeostasis. Prog Lipid Res. 2013. 52:249–276. PMID: 23500459

6. Parsons JB, and Rock CO. Its bacterial fatty acid synthesis a valid target for antibacterial drug discovery? Curr Opinion Microbiol. 2011; 14:544–549. PMID: 21862391

7. Sun, Y, Wilkinson BJ, Standiford TJ, Akinbi HT, and O’Riordan MXD. Fatty acids regulate stress resistance and virulence factor production for Listeria monocytogenes. J Bacteriol. 2012; 194:5274–5284. PMID: 22843841

8. Porta A, Török Z, Horvath I, Franceschelli S, Vígh L, et al. Genetic modification of the Salmonella membrane physical state alters the pattern of heat shock response. J Bacteriol. 2010; 192:1988–1998. PMID: 20139186

9. Schleifer KH, and Kroppenstedt RM. Chemical and molecular classification of staphylococci. Soc Appl Bacteriol Symp. Ser. 1990; 19:9S–24S. PMID: 211906

10. Singh VK, Hattangady DS, Giotis ES, Singh AK, Chamberlain NR, et al. Insertional inactivation of branched-chain α-keto acid dehydrogenase in Staphylococcus aureus leads to decreased branched-chain membrane fatty acid content and increased susceptibility to certain stresses. Appl Environ Microbiol. 2008; 74: 5882–5890. PMID: 18689519

11. Wilkinson BJ. Biology. In: K. B. Crossley KB, and Archer GL editors. The staphylococci in human disease. New York, NY. 1997. pp 1–38.

12. O’ Leary WM and Wilkinson SG. Gram positive bacteria. In: Ratledge C, Wilkinson SG, editors. Microbial Lipids Volume 1. London: Academic Press; 1988. pp. 117–201.

13. Kaneda T. Iso-and anteiso-fatty acids in bacteria: biosynthesis, function, and taxonomic significance. Microbiol Rev. 1991; 55: 288–302. PMID: 1886522

14. Legendre S, Letellier L, and Shechter E. Influence of lipids with branched-chain fatty acids on the physical, morphological and functional properties of Escherichia coli cytoplasmic membrane. Biochim Biophys Acta. 1980; 602:491–505. PMID: 6776984

15. Willecke K, and Pardee AB. Fatty acid-requiring mutant of Bacillus subtilis defective in branched chain alpha-keto acid dehydrogenase. J Biol Chem. 1971; 246:5264–5272. PMID: 4999353

16. Pelz A, Wieland KP, Putzbach K, Hentschel P, Albert K, et al. Structure and biosynthesis of staphyloxanthin from Staphylococcus aureus. J Biol Chem. 2005; 280: 32493–32498. PMID: 16020541

17. Wisniewska AJ, Widomska J, and Subczynski WK. Carotenoid membrane interactions in liposomes: effect of dipolar, monopolar, and non – polar carotenoids. Acta Biochem Pol. 2006; 53: 475–484. PMID: 16964324

18. Bramkamp M, and Lopez D. Exploring the existence of lipid rafts in bacteria. Microbiol Mol Biol Rev. 2015; 79:81–100. PMID: 25652542

19. Clauditz A, Resch A, Wieland KP, Peschel A, and Gotz F. Staphyloxanthin plays a role in the fitness of Staphylococcus aureus and its ability to cope with oxidative stress. Infect Immun. 2006; 74: 4950–4953. PMID: 16861688

20. Liu GY, Essex A, Buchanan JT, Datta V, Hoffman HM, et al. Staphylococcus aureus gold pigment impairs neutrophil killing and promotes virulence through its antioxidant activity. J Exp Med. 2005; 202: 209–215. PMID: 16009720

21. Liu CI, Liu GY, Song Y, Yin F, Hensler ME, et al. Cholesterol biosynthesis inhibitor blocks Staphylococcus aureus virulence. Science. 2008; 319: 1391–1394. PMID: 18276850

22. Campbell J, Singh AK, Santa Maria Jr JP, Kim Y, Brown S, et al. Synthetic lethalcompound combinations reveal a fundamental connection between wall teichoic acid andpeptidoglycan biosyntheses in Staphylococcus aureus. ACS Chem. Biol. 2011; 6:106–16.PMID: 20961110

23. Muthaiyan A, Silverman JA, Jayaswal RK, and Wilkinson BJ. Transcriptional profilingreveals that daptomycin induces the Staphylococcus aureus cell wall stress stimulon andgenes responsive to membrane depolarization. Antimicrob Agents Chemother. 2008; 52:980–990. PMID: 18086846

24. Song Y, Lunde CS, Benton BM and Wilkinson BJ. Studies on the mechanism of telavancin decreased susceptibility in a laboratory–derived mutant. Microbial Drug Res. 2013; 19: 247–255. PMID: 23551248

25. Clinical Laboratory Standards Institute. Methods for dilution antimicrobial susceptibility testing for bacteria that grow aerobically; approved standard, 7th ed. M7.A7. Clinical and Laboratory Standards Institute, Wayne, PA. 2006.

26. Zhu K, Bayles DO, Xiong A, Jayaswal RK, and Wilkinson BJ. Precursor and temperature modulation of fatty acid composition and growth of Listeria monocytogenes cold-sensitive mutants with transposon-interrupted branched-chain alpha-keto acid dehydrogenase. Microbiology. 2005; 151:615–623. PMID: 15699210.

27. Ray B, Ballal A, and Manna AC. Transcriptional variation of regulatory and virulence genes due to different media in Staphylococcus aureus. Microb Pathog. 2009; 47:94–100. PMID: 19450677

28. Oogai Y, Matsuo M, Hashimoto M, Kato F, Sugai M, et al. Expression of virulence factors by Staphylococcus aureus grown in serum. Appl Environ Microbiol. 2011; 77:8097–105. PMID: 21926198

29. Missiakas D, and Schneewind O. Growth and laboratory maintenance of Staphylococcus aureus. Current Protocols in Microbiol. 2013; DOI: 10.1002/9780471729259. PMID: 2340813

30. Yao J, and Rock CO. How bacterial pathogens eat host lipids: implications for the development of fatty acid synthesis therapeutics. J Biol Chem. 2015; doi: 10.1074/jbc.R114.636241

31. Chambers HF. The changing epidemiology of Staphylococcus aureus? Emerg Infect Dis. 2001; 7:178–182. PMID: 11294701

32. Moran GJ, A. Krishnadasan A, Gorwitz RJ, Fosheim GE, McDougal LK, et al. Methicillin-resistant S. aureus infections among patients in the emergency department. N Engl J Med. 2006; 355:666–674.

33. Fey PD, Endres JL, Yajjala VK, Yajjala K, Widhelm TJ, et al. A genetic resource for rapid and comprehensive phenotype screening of nonessential Staphylococcus aureus genes. MBio. 2012; 12: e00537–12. PMID: 23404398

34. Novick RP. Genetic systems in staphylococci. Methods Enzymol. 1991; 204: 587–636. PMID: 1658572

35. Davis AO, O’Leary JO, Muthaiyan A, et al. Characterization of Staphylococcus aureus mutants expressing reduced susceptibility to common house-cleaners. J Appl Microbiol. 2005; 98:364–72. PMID:15659191

36. Mishra NN, Liu GY, Yeaman MR, Nast CC, Proctor RA, et al. Carotenoid-related alteration of cell membrane fluidity impacts Staphylococcus aureus susceptibility to host defense peptides. Antimicrob Agents Chemother. 2011; 55:526–531. PMID: 21115796

37. Annous BA, Becker LA, Bayles DO, Labeda DP, and Wilkinson BJ. Critical role of anteiso-C_15:0_ fatty acid in the growth of Listeria monocytogenes at low temperatures. Appl Environ Microbiol. 1997; 63:3887–3894.

38. Edgcomb MR, Sirimanne S, Wilkinson BJ, Drouin P, and Morse RP. Electron paramagnetic resonance studies of the membrane fluidity of the foodborne pathogenic psychrotroph Listeria monocytogenes. Biochim Biophys Acta. 2000; 1463:31–42.

39. Parsons JB, Yao J, Frank MW, Jackson P, and Rock CO. Membrane disruption by antimicrobial fatty acids releases low molecular weight proteins from Staphylococcus aureus. J Bacteriol. 2012; 194: 5294–5304. PMID: 22843840

40. Townsend, D.E., and B.J. Wilkinson. 1992. Proline transport in Staphylococcus aureus: a high-affinity system and a low-affinity system involved in osmoregulation. J Bacteriol. 174:2702–2710. PMID: 1556088

41. Parsons J, Frank M, Jackson P, Subramanian C, and Rock CO. Incorporation of extracellular fatty acids by a fatty acid kinase-dependent pathway in Staphylococcus aureus. Mol Microbiol. 2014; 92: 234–245.

42. Parsons JB, Frank MW, Subramanian C, Saenkham P, and Rock CO. Metabolic basis for the differential susceptibility of Gram – positive pathogens for fatty acid synthesis inhibitors. Proc Natl Acad Sci. USA. 2011; 108: 15378–15383. PMID: 21876172

43. Payne JW, Gilvarg C. Size restriction on peptide utilization in Escherichia coli. J Biol Chem.1968; 243(23):6291–9. PMID: 26629334

44. Amezaga MR, Davidson I, McLaggan D, Verheul A, Abee T, Booth IR. The role of peptide metabolism in the growth of Listeria monocytogenes ATCC 23074 at high osmolarity. Microbiology. 1995; 141:41–49. PMCID: PMC167277

45. Hiron A, Borezee-Durant E, Piard JC, Juillard V. Only one of four oligopeptide transport systems mediates nitrogen nutrition in Staphylococcus aureus. J Bacteriol. 2007; 189:5119–5129.10.1128/JB.00274-07. PMID:17496096

46. Kaiser J.C., Sen S., Wilkinson B.J., Henrichs D. E. BrnQ1 in Staphylococcus aureus is a Leu/Val transporter required for determining branched-chain membrane fatty acids content. Abstr. Annu. Mtg. Am. Soc. Microbiol. 2016.

47. Qiu, X, Choudhry AE, Janson CA, Grooms M, Daines RA, et al. X Crystal structure and substrate specificity of ?-ketoacyl carrier protein synthase III (FabH) from Staphylococcus aureus. Protein Sci. 2007; 14: 2087–2094. PMID: 15987898

48. Altenbern RA. Cerulenin-inhibited cells of Staphylococcus aureus resume growth when supplemented with either a saturated or an unsaturated fatty acid. Antimicrob Agents Chemother. 1977; 11: 574–576PMID: 856007

49. Holman R, Adams C, Nelson R, Grater S, Jaskiewicz J, et al. Patients with anorexia nervosa demonstrate deficiencies of selected essential fatty acids, compensatory changes in nonessential fatty acids and decreased fluidity of plasma lipids. J Nutr. 1995; 125:901–907. PMID: 7722693.

50. Nakamura T, Azuma A, Kuribayashi T, Sugihara H, Okuda S, et al. Serum fatty acid levels, dietary style and coronary heart disease in three neighboring areas in Japan: the Kumihama study. Br J Nutr. 2003; 89:267–272. PMID: 12575911

51. Shimomura, Y, Sugiyama S, Takamura T, Kondo T, and Ozawa T. Quantitative determination of the fatty acid composition of human serum lipids by high-performance liquid chromatography. J Chromatogr. 1986; 383:9–17. PMID: 3818849

52. Psychogios N, Hau DD, Peng J, Guo AC, Mandal AC, et al. The human serum metabolome. PLoS ONE. 2011; 6(2): e16957. PMID: 21359215

53. Brinster S, Lamberet G, Staels B, Trieu-Cuot P, Gruss A, et al. Type II fatty acid synthesis is not a suitable antibiotic target for Gram-positive pathogens. Nature. 2009; 458: 83–86. PMID: 19262672

54. Clarke SR, Mohamed R, Bian L, Routh AF, Kokai-Kun JF, Mond JJ, et al. The Staphylococcus aureus surface protein IsdA mediates resistance to innate defenses of human skin. Cell Host Microbe. 2007; 1:199–212. doi: 10.1016/j.chom.2007.04.005 PMID:18005699

55. Stewart ME. Sebaceous gland lipids. Semin Dermatol. 1992; 11:100–105.PMID:1498012

56. Hamosh M. Protective function of proteins and lipids in human milk. Biol Neonate. 1998; 74:163–176. PMID: 9691157

57. Takigawa H, Nakagawa H, Kuzukawa M, Mori H, Imokawa G. Deficient production of hexadecenoic acid in the skin is associated in part with the vulnerability of atopic dermatitis patients to colonization by Staphylococcus aureus. Dermatology. 2005; 211:240–248. 10.1159/000087018. PMID: 16205069

58. Cartron ML, England SR, Chiriac AI, Josten M, Turner R, et al. Bactericidal activity ofthe human skin fatty acid cis-6-hexadecanoic acid on Staphylococcus aureus. Antimicrob Agents Chemother. 2014; 58:3599–3609. PMID: 24709265

59. Kenny JG, Ward D, Josefsson E, Jonsson IM, Hinds J, Lindsay JA, et al. The Staphylococcus aureus response to unsaturated long chain free fatty acids: survival mechanisms and virulence implications. PLOS One. 2009. DOI: 10.1371/journal.pone.0004344

60. Bose, JL, Daly SM, Hall RR, and Bayles KW. Identification of the vfrAB operon in Staphylococcus aureus: A novel virulence factor regulatory locus. Infect Immun. 2014; 82: 1813–1822.

61. Sabirova JS, Hernalsteens JP, De Backer S, et al. Fatty acid kinase A is an important determinant of biofilm formation in Staphylococcus aureus USA300. BMC Genomics. 2015; 16:861. Doi: 10.1186/s12864-015-1956-8. PMID: 26502874

62. Stoll H, Dengjel J, Nerz C & Götz F. Staphylococcus aureus deficient in lipidation of prelipoproteins is attenuated in growth and immune activation. Infect Immun. 2005; 73: 2411–2423. PMCID: PMC1087423

63. Nguyen MH, Hanzelmann D, Härtner T, Peschel A, Götz F. Skin-specific unsaturated fatty acid boost Staphylococcus aureus innate immune response. Infect Immun. 2015; doi:10.1128/IAI.00822-15

64. Silbert DF, Ladenson RC, and Honegger JL. The unsaturated fatty acid requirement in Escherichia coli. Temperature dependence and total replacement by branched-chain fatty acids. Biochim Biophys Acta. 1973; 311: 349–361. PMID: 4580982

65. Parsons JB, Frank MW, Rosch JW, and Rock CO. Staphylococcus aureus fatty acid auxotrophs do not proliferate in mice. Antimicrob Agents Chemother. 2013; 57: 5729–5732. PMID: 23979734.

66. Malachowa N. Whitney AR, Kobayashi SD, Sturdevant DE, Kennedy AD, et al. Global changes in Staphylococcus aureus gene expression in human blood. PLoS ONE. 2011; 6:e18617. PMID: 21525981

67. Garber ED. The host as a growth medium. Annals of the New York Academy of Sciences.1960; 88:1187–1194. doi: 10.1111/j.1749-6632.

68. Brown SA, Palmer KL, Whiteley M. Revisiting the host as a growth medium. Nat Rev Microbiol. 2008; 6: 657–666. doi: 10.1038/nrmicro1955

69. Massey RC, Dissanayeke SR, Cameron B, Ferguson D, Foster TJ, and Peacock SJ. Functional blocking of Staphylococcus aureus adhesins following growth in ex vivo media. Infect Immun. 2002; 70:5339–5345. PMID: 12228257

70. Krismer B, Liebeke M, Janek D, Nega M, Rautenberg M, Hornig G, et al. Nutrient limitation governs Staphylococcus aureus metabolism and niche adaptation in the human nose. PLoS Pathogens. 2014;10: e1003862 doi: 10.1371/journal.ppat.1003862

71. Citterio L, Franzyk H, Palarasah Y, Andersen TE, Mateiu RV, Gram L. Improved in vitro evaluation of novel antimicrobials: potential synergy between human plasma and antibacterial peptidomimetics, AMPs and antibiotics against human pathogenic bacteria. Research in Microbiology. 2016: 167: 72–82., 10.1016/j.resmic.2015.10.002

